# High density SNP array and reanalysis of genome sequencing uncovers CNVs associated with neurodevelopmental disorders in KOLF2.1J iPSCs

**DOI:** 10.1101/2023.06.26.546614

**Authors:** Carolina Gracia-Diaz, Jonathan E. Perdomo, Munir E. Khan, Brianna Disanza, Gregory G. Cajka, Sunyimeng Lei, Alyssa Gagne, Jean Ann Maguire, Thomas Roule, Ophir Shalem, Elizabeth J. Bhoj, Rebecca C. Ahrens-Nicklas, Deborah French, Ethan M. Goldberg, Kai Wang, Joseph Glessner, Naiara Akizu

**Affiliations:** Raymond G. Perelman Center for Cellular and Molecular Therapeutics, The Children’s Hospital of Philadelphia, Philadelphia, PA, USA; Department of Pathology and Laboratory Medicine, University of Pennsylvania, Philadelphia, PA, USA; School of Biomedical Engineering, Drexel University, Philadelphia, PA 19104, USA; Center for Applied Genomics, The Children’s Hospital of Philadelphia, Philadelphia, PA, USA; Department of Pediatrics, University of Pennsylvania, Philadelphia, PA, USA; Department of Genetics, University of Pennsylvania, Philadelphia, PA, USA; Departmen of Neurology, University of Pennsylvania, Philadelphia, PA, USA; Division of Neurology, Department of Pediatrics, The Children’s Hospital of Philadelphia, Philadelphia, PA, USA

**Author notes:** Correspondance (K.W.), (J.G.), (N.A.).

## Abstract

The KOLF2.1J iPSC line was recently proposed as a reference iPSC to promote the standardization of research studies in the stem cell field. Due to overall good performance differentiating to neural cell lineages, high gene editing efficiency, and absence of genetic variants associated to neurological disorders KOLF2.1J iPSC line was particularly recommended for neurodegenerative disease modeling. However, our work uncovers that KOLF2.1J hPSCs carry heterozygous small copy number variants (CNVs) that cause *DTNBP1, JARID2* and *ASTN2* haploinsufficiencies, all of which are associated with neurological disorders. We further determine that these CNVs arose *in vitro* over the course of KOLF2.1J iPSC generation from a healthy donor-derived KOLF2 iPSC line and affect the expression of DNTBP1, JARID2 and ASTN2 proteins in KOLF2.1J iPSCs and neural progenitors. Therefore, our study suggests that KOLF2.1J iPSCs carry genetic variants that may be deleterious for neural cell lineages. This data is essential for a careful interpretation of neural cell studies derived from KOLF2.1J iPSCs and highlights the need for a catalogue of iPSC lines that includes a comprehensive genome characterization analysis.

## Introduction

The derivation of the first human embryonic stem cells (hESC)^1^ followed by the generation of induced pluripotent stem cells (iPSC) from somatic cells^2^ and more recent advances in genome engineering methods^3^ have transformed the way we study human biology and diseases. Today, we can generate virtually every human cell type *in vitro* and we can do so in the genetic background of choice. This progress has particularly benefited research focused on the study of human brain development, mechanisms and disorders by providing a nearly universal access to human neural cells, which are otherwise of limited accessibility.

Since the differentiation of the first neurons from hESC and iPSCs (collectively known as human pluripotent stem cells, hPSCs)^4, 5^, protocols for 2D and 3D neural tissue generation have exponentially increased and expanded the horizon for mechanistic and therapeutic discoveries of human brain disorders. Yet, these advances come with new challenges for the field, including the multiple sources of technical and biological variables that lead to the expression of phenotypes often difficult to reconcile^6, 7^. To overcome these challenges, early efforts focused on improving the reproducibility and efficiency of neural differentiation protocols^8–11^. However, even with the most reproducible protocol, the genetic background of an individual hPSC line can significantly contribute to the phenotypic heterogeneity, just like in the general population. Studies using high quality genome assemblies obtained with long read sequencing estimate that each human genome contains approximately ∼4,000,000 single nucleotide variants (SNVs), ∼800,000 insertion deletions (indels) and 25,000 structural variants (SV), such as copy number variants (CNVs) or inversions and insertions of >50bp^12^. Given that recapitulating the large genetic heterogeneity of the human population in a dish would limit the identification of significant genotype-phenotypes relations, recent efforts have proposed to homogenize the genetic background by adopting a widespread use of reference hPSCs^13^. The success of this idea is supported by decades of key discoveries in isogenic strains of model organisms^14, 15^.

In a recent effort to generate a readily available collection of isogenic iPSC carrying mutations associated with neurodegenerative disorders for distribution and collaboration across different laboratories, the iPSC Neurodegenerative Disease Initiative (iNDI) sought to identify an overall well-performing iPSC line that would serve as a reference for the research community^13, 16^. To achieve this goal, nine iPSCs lines from public repositories were selected for a deep functional and genomic characterization. One of these iPSC lines, named KOLF2.1J, was generated from the KOLF2_C1 iPSC line by CRISPR/Cas9 correction of a heterozygous 19bp deletion in *ARID2*^17^, a variant likely pathogenic for a neurodevelopmental disorder known as Coffin-Siris syndrome. After a deep characterization of several subclones of the nine iPSC lines, KOLF2.1J was selected as a candidate reference iPSC line for its good performance at pluripotent stage and differentiation to neural cell linages, high CRISPR-Cas9 editing efficiency and low burden of genetic variants associated with neurological disorders^13^. To further test KOLF2.1J for the presence of genetic variants that might affect the experimental interpretations, additional genomic analyses were performed, including comparative studies with the parental fibroblasts (from a 55-59yo healthy male donor) and reprogrammed iPSC clone KOLF2-C1. These analyses revealed ∼3.3 million high confidence SNPs and indels affecting coding regions in KOLF2.1J. Twenty-five of these were present only in KOLF2.1J and not in KOLF-C1 iPSCs and 37 were inherited from the parental fibroblasts. Only three of these variants affected genes predicted to be intolerant to loss of function or listed as haploinsufficent (*COL3A1*, *SHOX* and *DEDD*) but none were suspected to compromise neurological disease modeling. Notably, structural variants of 50 bp-5 Mbp in size were not assessed in these genomic analyses due to technical limitations of the selected methods^13^.

Here we show that KOLF2.1J iPSCs carry at least 5 small CNVs (<1Mbp), two of which cause *DTNBP1, JARID2* and *ASTN2* haploinsufficiency. Given the association of these genes with neurodevelopmental disorders^18–27^ and constraints for disruptive variants in human genomes^28–30^ our work uncovers that KOLF2.1J iPSCs carry genetic variants that may affect the interpretation of developmental and neural cell phenotypes. Our work further highlights the need for the inclusion of structural variants in genome sequencing analyses of iPSC lines and the creation of a catalogue of reference iPSCs which shall include a comprehensive characterization of genes predicted to be intolerant to disruptive variation during human development.

## Results

### Genome integrity validation detects CNVs in KOLF2.1J iPSCs

Owing to their deep characterization and malleability for genome editing, we recently adopted KOLF2.1J iPSCs (The Jackson Laboratory, JIPSC1000) for neurodegenerative and neurodevelopmental studies conducted in our laboratories. Our projects originate from the identification of candidate pathogenic variants through exome and genome sequencing of children affected with progressive or non-progressive neurodevelopmental disorders. We then design *in silico* and biochemical studies to test the impact of the variants on the encoded proteins and functionally validate those with a significant molecular effect. The hPSC based neural differentiation models constitute the top tier of our studies for which we reprogram patient cells or use CRISPR-Cas9 to introduce the candidate variants in an isogenic background to compare neural cell phenotypes between variants and controls.

Following this strategy, we selected several variants to CRISPR-Cas9 edit in KOLF2.1J iPSCs and attest their high editing efficiency. After gene editing and expansion of edited KOLF2.1J clones to generate frozen stocks, we applied our routine quality control tests, including a genome integrity validation. For this purpose, we extracted genomic DNA (gDNA) from the edited KOLF2.1J iPSCs and subsequently analyzed with an array containing common and rare alleles of ∼650.000 single nucleotide polymorphisms (SNPs) spanning the whole human genome. As a method for structural integrity validation we analyzed the array for CNVs causing deletions or duplications of genome fragments using PennCNV^31^. Furthermore, to rule out the possibility of undesired genetic events arising during the editing process, we compared the results with the CNV analysis of an unedited KOLF2.1J iPSC clone. Although, the comparison confirmed absence of acquired CNVs in most of the edited clones, results unexpectedly revealed recurrent small CNVs (<1Mbp) in all edited and unedited KOLF2.1J iPSCs (Table 1), suggesting the presence of these CNVs in our KOLF2.1J iPSC stock. To corroborate this finding, we conducted the same SNP array based CNV analysis in one of our KOLF2.1J iPSCs stocks at +1 passage (p3), KOLF2.1J iPSCs carrying the doxycycline inducible *NGN2* transgene (both kindly donated by Dr. Skarnes) and a KOLF2.1J stock at p2 recently received from The Jackson Laboratory. Results confirmed the presence of the 5 CNVs in all the stock KOLF2.1J iPSCs (Table 1, Fig 1 and Supplementary Fig 1 and 2).

**Figure 1:**
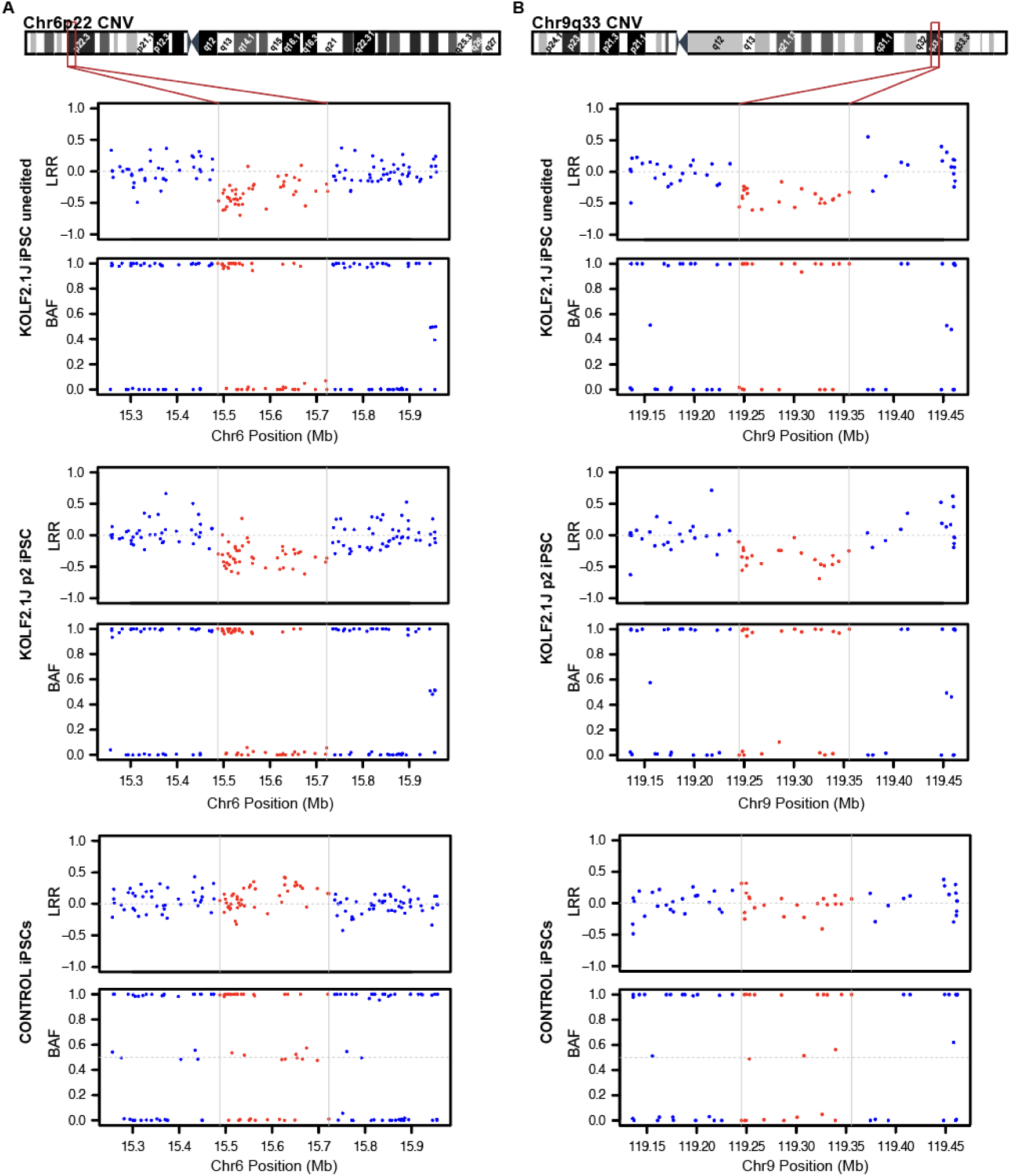
High density SNP array uncovers CNVs affecting coding genes in Chr6p22 and Chr9q33 in KOLF2.1J iPSCs. **(A)** Chromosome 6 cytoband schematics (top) and Log R Ratio (LRR) and B Allele frequency (BAF) plots (bottom) show reduction of signal intensity and a loss of heterozygosity in 6p22 region of unedited and p2 KOLF2.1J iPSCs compared to a control iPSC line. (**B)** Chromosome 9 cytoband schematics (top) and LRR and BAF plots (bottom) show reduction of signal intensity and a loss of heterozygosity in 9q33 region of KOLF2.1J iPSCs unedited and at p2 compared to a control iPSC line.

**Figure 2:**
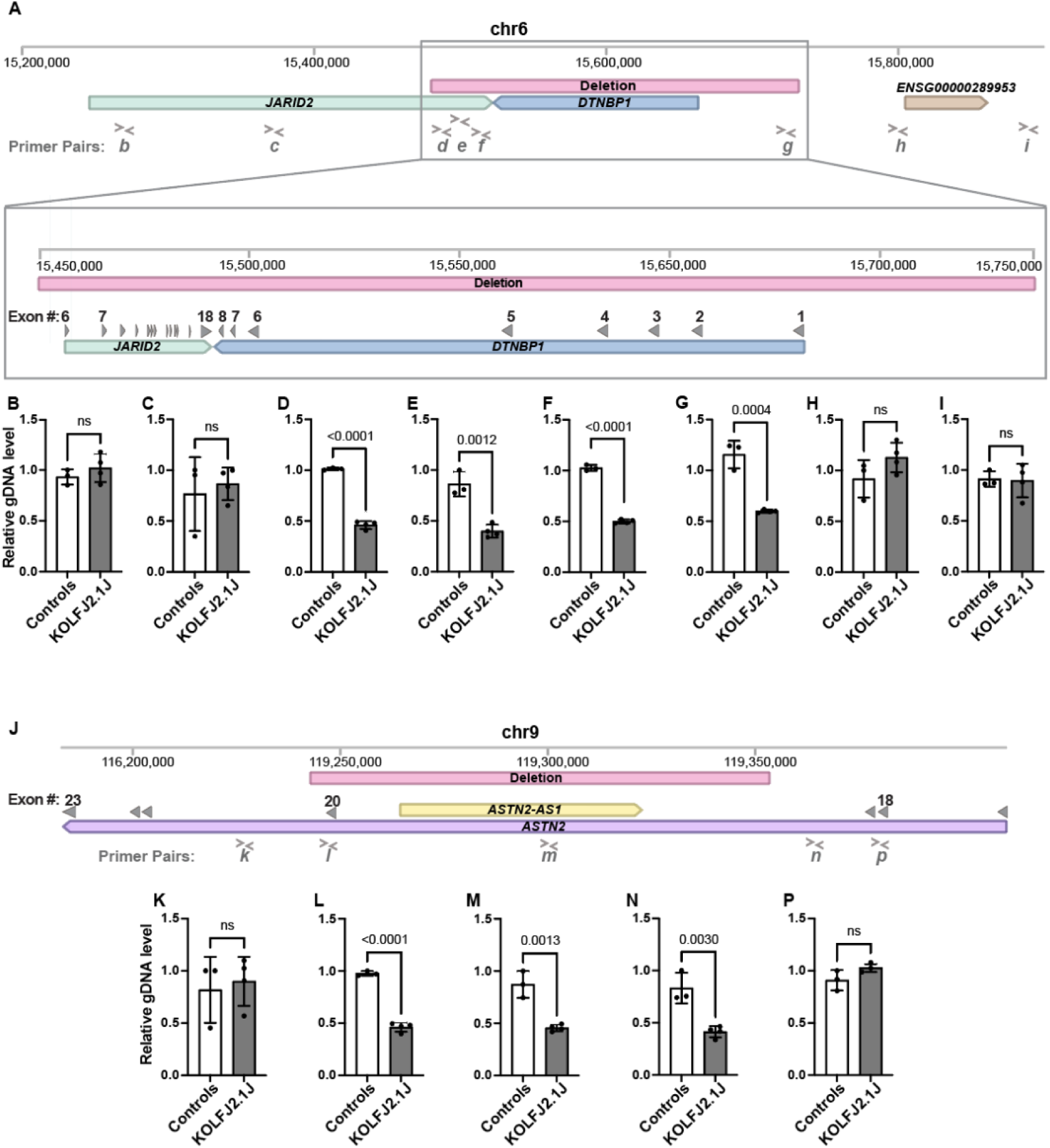
gDNA qPCR analysis confirms the presence of chr6p22 and chr9q33 CNV in KOLF2.1J iPSCs. **(A)** Schematic representation of Chromosome 6 (Chr6:15,200,000-15,900,000). Window illustrates the 0.2 Mb deleted region (red bar) and affected genes, *JARID2* (green bar) and *DTBNP1* (blue bar). (**B-I)** Genomic DNA qPCR results showing half levels of amplification within the Chr6 deleted CNV region in KOLF2.1J compared to control iPSCs (D-G) and regions upstream and downstream of the deletion (A-B and H-I). (**J)** Schematic representation of Chromosome 9 (Chr9:116,200,000-119,400,000) with the 0.1 Mb deleted region (red bar) and affected genes, *ASTN2* (purple bar) and *ASTN-AS1* (yellow bar). (**K-P)** Genomic DNA qPCR results showing half levels of amplification within the Chr9 deleted CNV region in KOLF2.1J compared to control iPSCs (L-N) and regions upstream and downstream of the deletion (K and P). Graphs show mean +/-SD of n=3 control hPSC lines and n=4 KOLF2.1J iPSC independent stocks. p-values were calculated with a two-sided unpaired t-test. ns=non-significant. Grey filled arrowheads indicate exons. > and < symbols indicate qPCR primer pair positions.

**Table 1:** CNVs detected in KOLF2.1J iPSC lines by high density SNP array.

| Sample | Cytoband | Aprox. lengh | Aprox. Coordinates | CNV type | Affected Genes |
| --- | --- | --- | --- | --- | --- |
| KOLF2.1J<br>Unedited | 3p13 | 0.1 Mb | Chr3:72,289,657-72,404,542 | Deletion | No coding genes |
|  | 3q14 | 0.3 Mb | Chr3:61,242,795-61,528,270 | Duplication | No coding genes |
|  | 6p22 | 0.2 Mb | Chr6:15,487,649-15,722,102 | Deletion | *JARID2* and *DTNBP1* |
|  | 9p22 | 0.1 Mb | Chr9:119,244,942-119,355,584 | Deletion | *ASTN2* and *ASTN2-AS1* |
|  | 18q22 | 0.1 Mb | Chr18:62,115,075-62,264,706 | Duplication | No coding genes |
| KOLF2.1J<br>Edited 1 | 3p13 | 0.1 Mb | Chr3:72,289,657-72,404,542 | Deletion | No coding genes |
|  | 3q14 | 0.3 Mb | Chr3:61,242,795-61,528,270 | Duplication | No coding genes |
|  | 6p22 | 0.2 Mb | Chr6:15,487,649-15,722,102 | Deletion | *JARID2* and *DTNBP1* |
|  | 9p22 | 0.1 Mb | Chr9:119,244,942-119,355,584 | Deletion | *ASTN2* and *ASTN2-AS1* |
|  | 18q22 | 0.1 Mb | Chr18:62,115,075-62,264,706 | Duplication | No coding genes |
| KOLF2.1J<br>p2 | 3p13 | 0.1 Mb | Chr3:72,289,657-72,404,542 | Deletion | No coding genes |
|  | 3q14 | 0.3 Mb | Chr3:61,242,795-61,528,270 | Duplication | No coding genes |
|  | 6p22 | 0.2 Mb | Chr6:15,487,649-15,722,102 | Deletion | *JARID2* and *DTNBP1* |
|  | 9p22 | 0.1 Mb | Chr9:119,244,942-119,355,584 | Deletion | *ASTN2* and *ASTN2-AS1* |
|  | 18q22 | 0.1 Mb | Chr18:62,115,075-62,264,706 | Duplication | No coding genes |
| KOLF2.1J<br>p3 | 3p13 | 0.1 Mb | Chr3:72,289,657-72,404,542 | Deletion | No coding genes |
|  | 3q14 | 0.3 Mb | Chr3:61,242,795-61,528,270 | Duplication | No coding genes |
|  | 6p22 | 0.2 Mb | Chr6:15,487,649-15,722,102 | Deletion | *JARID2* and *DTNBP1* |
|  | 9p22 | 0.1 Mb | Chr9:119,244,942-119,355,584 | Deletion | *ASTN2* and *ASTN2-AS1* |
|  | 18q22 | 0.1 Mb | Chr18:62,115,075-62,264,706 | Duplication | No coding genes |
| KOLF2.1J-NGN2 p10 | 3p13 | 0.1 Mb | Chr3:72,289,657-72,404,542 | Deletion | No coding genes |
|  | 3q14 | 0.3 Mb | Chr3:61,242,795-61,528,270 | Duplication | No coding genes |
|  | 6p22 | 0.2 Mb | Chr6:15,487,649-15,722,102 | Deletion | *JARID2* and *DTNBP1* |
|  | 9p22 | 0.1 Mb | Chr9:119,244,942-119,355,584 | Deletion | *ASTN2* and *ASTN2-AS1* |
|  | 18q22 | 0.1 Mb | Chr18:62,115,075-62,264,706 | Duplication | No coding genes |

### 6p22 and 9p33 deletion CNVs are associated with neurodevelopmental disorders

With an estimate of ∼25,000 structural variants (SV) carried per genome in average, it is not surprising that most hPSCs used for research also carry SV^32^. Furthermore, numerous studies have demonstrated that the origin of the parental cell, the reprogramming process and the culture conditions can contribute to structural genetic variation^33–38^. KOLF2.1J iPSCs were derived from a CRISPR-Cas9 mediated correction of a 19 bp deletion in one copy of *ARID2* present in the parental KOLF2-C1 iPSC line and expanded to hundreds of replicate vials to ensure their availability for distribution^13^. Although all these procedures are a known source of genetic structural variation^36, 38, 39^, most type of structural variants were not reported in KOLF2.1J iPSCs^13^. Given the discrepancy between these and our results, we took a closer look to the SNPs at the CNV regions of KOLF2.1J iPSCs by visualizing their signal intensity (Log R Ratio (LRR)) and B allele frequency (BAF). Consistent with a heterozygous deletion, we observed a drop in LRR values of SNPs located within 3p13, 6p22 and 9q33 CNV regions relative to those in adjacent diploid regions, while the signal intensity of duplicated CNV regions in 3p14 and 18q33 was slightly above (Fig 1, Supplementary Fig 1 and 2). Furthermore, as expected by one copy loss of a DNA fragment, the regions with reduced LRRs also showed lack of heterozygous SNPs in KOLF2.1J iPSCs (Fig 1, Supplementary Fig 1 and 2) while the control iPSC line we analyzed in parallel showed LRR and BAF values of 6p22, 9q33 and adjacent SNPs that were consistent with diploid state (Fig 1).

Among the 5 CNVs identified, two span coding regions. One is a ∼0.2Mbp heterozygous copy loss located in chromosome 6p22 which deletes one copy of the entire *DTNBP1* gene and 11 out of the 18 coding exons of *JARID2* (Fig 2A). *DTNBP1* encodes Dysbindin, a protein involved in lysosome-related organelle biogenesis which expression increases with neuronal differentiation in the developing and adult human brain (Supplementary Fig 3A)^40, 41^. While there is no evidence of pathogenic consequences for heterozygous *DTNBP1* loss, homozygous nonsense mutations cause Hermansky-Pudlak syndrome 7 (OMIM #614076)^42–44^ and the locus is associated with increased susceptibility for schizophrenia^25–27^. More concerning for studies involving neural cell types is the heterozygous loss of *JARID2*, a gene highly intolerant to loss of function (pLI=1, pLOF o/e=0.09)^28^, and copy number variation (pHaplo=0.99, pTriplo=0.99 and pHI=0.59)^29, 30^ in the human population (Table 2). *JARID2* is highly expressed in pluripotent cells and in the developing human brain with expression slightly declining as neurons differentiate (Supplementary Fig 3B)^40, 41^. As part of the Polycomb Repressive Complex 2 (PRC2) regulatory subunit, JARID2 is critical for embryonic development and tissue homeostasis^45^. Notably, similar CNVs causing partial or full deletion of one copy of *JARID2*, are the molecular basis for a neurodevelopmental delay with variable intellectual disability and dysmorphic facies syndrome (OMIM #620098)^18–21^.

**Table 2:** Constriction metrics and OMIM phenotypes of genes affected by KOLF2.1J CNVs compared to *ARID2*.

| Gene | OMIM # | pHaplo<sup>29</sup> | pTriplo<sup>29</sup> | Decipher p(HI)<sup>30</sup> | GenomAD<sup>28</sup><br>pLI | GnomAD<sup>28</sup><br>pLOF o/e |
| --- | --- | --- | --- | --- | --- | --- |
| *JARID2* | 620098 | 0.997 | 0.992 | 0.58982626\|12.14% | 1 | 0.09 (0.05 - 0.19) |
| *DTNBP1* | 614076 | 0.769 | 0.272 | 0.148024092\|43.28% | 0 | 0.59 (0.37 - 0.98) |
| *ASTN2* |  | 0.884 | 0.672 | 0.952118309\|2.19% | 1 | 0.14 (0.08 - 0.25) |
| *ARID2* | 617808 | 0.997 | 0.981 | 0.622938783\|11.02% | 1 | 0.04 (0.02 - 0.1) |

The other CNV affecting a coding region is a ∼0.1Mbp heterozygous deletion in chromosome 9q33 overlapping with the exon 20 of *ASTN2* and its antisense gene (*ASTN2-AS1*) (Fig 2J). *ASTN2* encodes a protein involved in trafficking and degradation of neuronal cell adhesion molecules important for the regulation of neuronal migration and synaptic development^46^. Similar to *JARID2*, *ASTN2* is a gene with low tolerance for loss of function and copy number variation (pLI=1, pHaplo=0.88, pTriplo=0.672 and pHI=0.95)^28–30^ (Table 2) and its genetic disruption is a risk factor for multiple neurodevelopmental disorders^22–24^. The expression of *ASTN2* is low at pluripotent stage but increases in neurons and is widely expressed in the developing and adult brain (Supplementary Fig 3C)^40, 41^. Although we have not experimentally investigated the consequence of this CNV for *ASTN2* expression, the loss of exon 20 is predicted to lead to a premature stop codon and nonsense mediated RNA decay.

To further validate the SNP array CNV results with an alternative method, we performed a quantitative PCR (qPCR) analysis of genomic DNA (gDNA) with a set of primers targeting the CNVs and upstream and downstream regions not predicted to be affected by the CNVs. As a positive control for sensitivity of this approach distinguishing one copy from two copies of genome fragments, we used a previously reported patient iPSC line carrying a heterozygous deletion within *EZH1* gene^47^. As expected, the level of amplification of *EZH1* region in this patient-derived iPSCs was half of the control hPSCs (Supplementary Fig 4). Similarly, the qPCR with primers targeting 6p22 and 9q33 CNVs showed half levels in KOLF2.1J iPSCs compared to control hPSCs, while the level of amplification of diploid neighboring regions were similar between all the hPSC lines (Fig 2B-I and K-P). These data confirm *JARID2* and *ASTN2* haploinsufficiency in several stocks of KOLF2.1J iPSCs that we independently received.

### Reanalysis of genome sequencing data confirms that KOLF2.1J iPSCs carry CNVs associated with neurodevelopmental disorders

Given that our data shows that the stock of KOLF2.1J iPSCs being currently distributed carry CNVs associated with neurodevelopmental disorders, we wondered whether the 5 CNVs were already present in the KOLF2.1J genome sequencing data, yet missed during the analysis previously reported^13^. In support of this possibility, we found that the original structural analysis of the genome sequencing was performed with an algorithm that considers discordant read-pairs and split-reads for CNV calling^48^, which can lead to false negatives when there is not enough evidence of aberrant read pairs passing quality control filtering.

The detection of structural variants from short read genome sequencing studies remains challenging. However, several algorithms that consider read depth in their analysis show improved sensitivity for CNV calling from short-read genome sequencing data^49^. To determine if the genetic background of KOLF2.1J iPSCs includes the 5 CNVs identified in our stocks, we downloaded the genome sequencing data deposited in the Alzheimer’s Disease Workbench (https://doi.org/10.34688/KOLF2.1J.2021.12.14)13 and aligned to the reference genome in paired-end short read mode. Then, we calculated the Log2 Ratio and BAF across alignments with read coverages and allele counts obtained for each CNV region using an in-house developed software. Strikingly, the five CNVs identified by the SNP array were also present in the genome sequencing data of KOLF2.1J iPSCs (Fig 3 and Supplementary Fig 5). In addition, the reanalysis allowed us to define the breakpoints of each CNV at base-resolution, which revealed that the 3p14 duplication is larger than indicated by the SNP array CNV calling (Supplementary Fig 5B). Notably, these new coordinates indicate that the CNV 3p14 duplication overlaps with coding regions of *FHIT* and *PTPRB,* although functional consequences are uncertain because it is difficult to tell whether there is a tandem duplication or an extra copy elsewhere in the genome.

**Figure 3:**
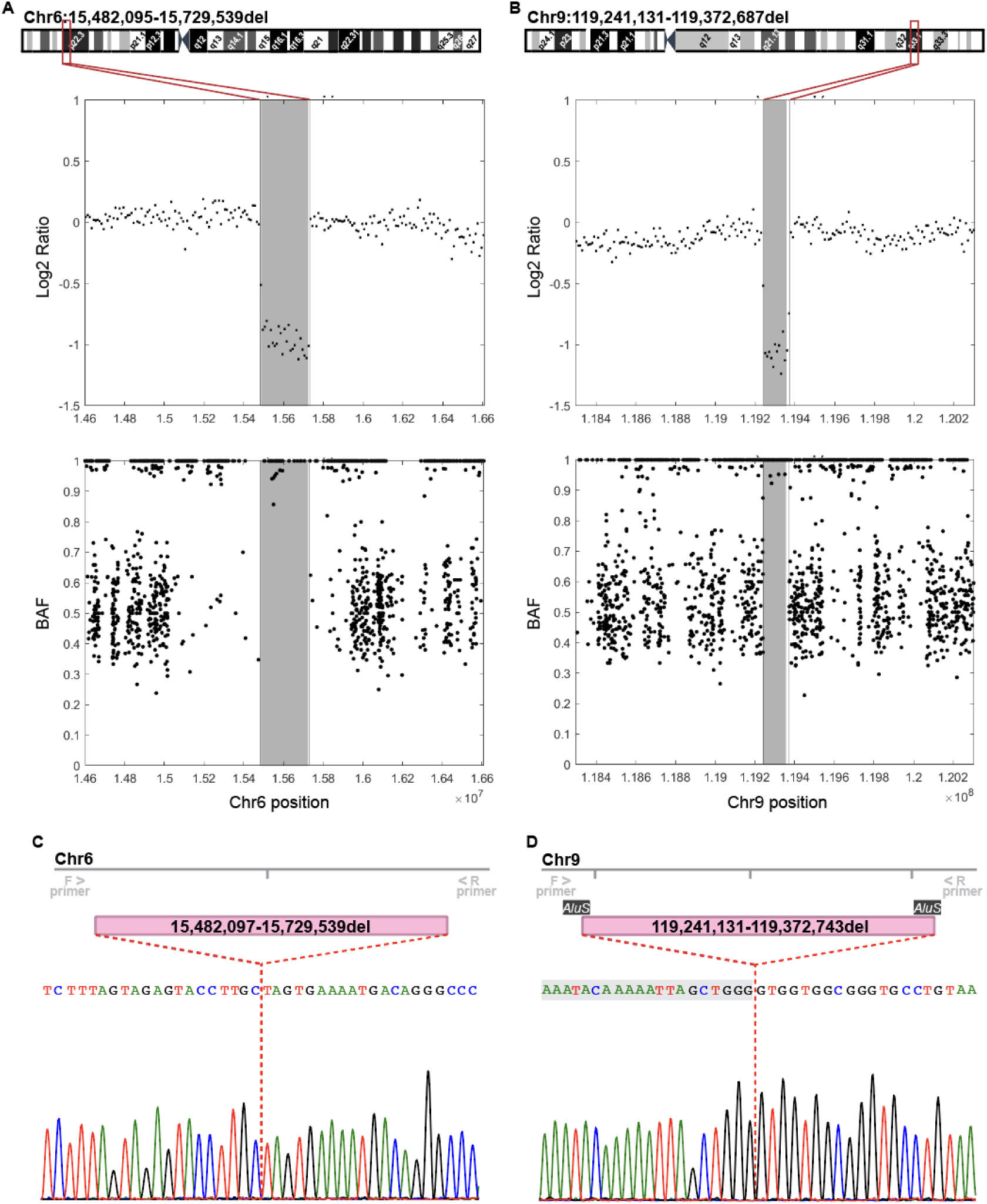
Genome sequencing reanalysis confirms CNVs at Chr6p22 and Chr9q33. **(A, B)** Log2 ratio and BAF plots show reduction of signal intensity (Log2 ratio) and a loss of heterozygosity (BAF) in Chr6p22 (A) and Chr9q33 (B) CNV regions compared to diploid flanking regions. Shadowed area represents the CNV defined by the SNP array and vertical lines the base-resolution breakpoints determined from the genome sequencing data. Chromosome 6 and 9 cytoband schematics with the CNV breakpoints at base resolution are shown on the top. **(C, D)** Sanger sequencing of PCR amplicons including ∼200bp up-and downstream of the predicted breakpoints confirm deletion CNVs in Chr6(C) and Chr9 (D) and adjust the breakpoint positions. Nucleotides shadowed in grey (D) are part of the 56 bp long microhomology domain of two *AluS* repetitive elements localized in Chr9 CNV breakpoints.

To further validate the genome sequencing CNV analysis results, we analyzed the Chr6 and Chr9 CNV breakpoint junctions in KOLF2.1J iPSCs by Sanger sequencing. Results confirmed the breakpoint coordinates for the Chr6 CNV, with only one bp discrepancy from the genome sequencing analysis (Fig 3C). For the Chr9 CNV, the Sanger sequencing result corrected the breakpoint junction by 56bps (Fig 3D). Interestingly, we noted that these 56bps were repeated almost identically (except for two nucleotides) in the sequences surrounding the 5’ and 3’ breakpoints of the Chr9 CNV. Thus, we had a closer look at these genomic regions and uncovered that the 56bps repeats are part of the microhomologies of two *AluS* elements flanking the Chr9 CNV. This finding indicates that the Chr9 CNV arose from an *Alu/Alu* mediated genomic rearrangement, which is a known source of genomic structural variation and estimated to cause ∼0.3% of human genetic diseases^50^.

### Chr6 and Chr9 CNVs arose *in vitro* and affect the expression of *JARID2*, *DTNBP1* and *ASTN2* genes in KOLF2.1J iPSC line

Having confirmed that KOLF2.1J iPSCs carry CNVs that are associated with neurological disorders, we next wondered whether these CNVs were already present in the healthy, 55-59 year-old donor. This possibility would suggest that the CNVs are not deleterious. Of note, KOLF2.1J iPSC line was derived from a subclone of KOLF2 iPSCs reprogramed from the donor’s fibroblasts, and KOLF2 genomic data is publicly accessible through the European Nucleotide Archive. Therefore, we downloaded the available KOLF2 SNP genotyping data, which contains LRR and BAF information for 511,966 SNPs, and analyzed for CNVs. CNV calling revealed that only Chr3p14.2 and Chr18q22.1 duplication CNVs were present in KOLF2 iPSCs (Table 3). Furthermore, visual analysis of SNPs within the Chr3p13 showed BAFs of ∼0.3 and ∼0.7 which suggests the subclonal presence of this CNV (Table 3 and Supplementary Fig 6A-C). Strikingly, Chr6 and Chr9 CNVs were not called nor detected by visual review of SNPs LRRs and BAFs in KOLF2 iPSCs (Table 3 and Fig 4A,B), indicating that they arose *in vitro* over the course of culturing, subcloning and CRISPR/Cas9 editing from KOLF2 iPSC to KOLF2.1J iPSC generation.

**Figure 4:**
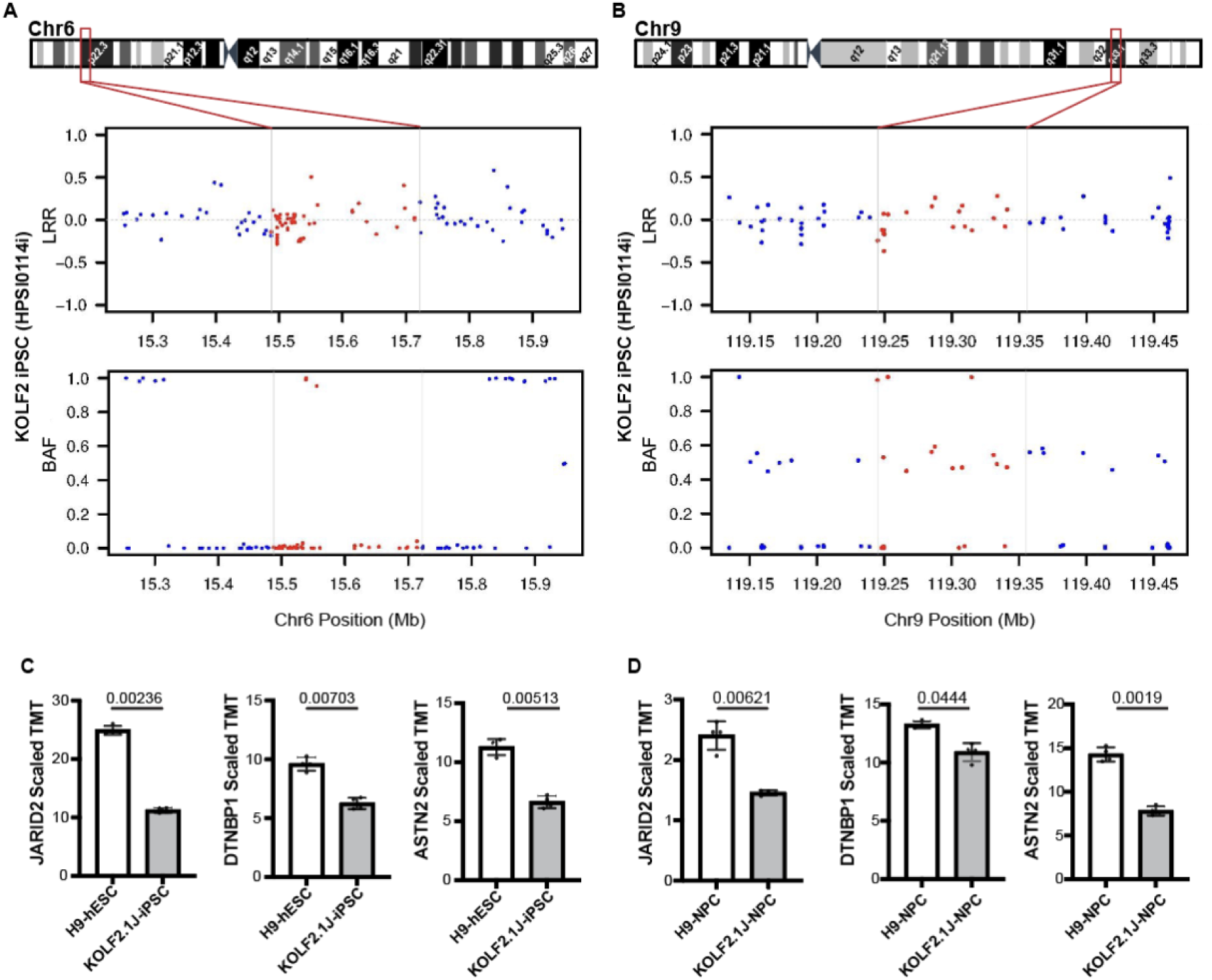
Chr6p22 and Chr9q33 CNVs arose *in vitro* and affect the expression of *JARID2*, *DTNBP1* and *ASTN2* genes. **(A, B)** LRR and BAF plots of the KOLF2 iPSC line SNP array show diploid Chr6 (A) and Chr9 (B) regions, indicating that the CNVs arose in culture between KOLF2 to KOLF2.1J generation. Horizontal lines and red dots depict SNPs that fall within KOLF2.1J CNV regions. **(C, D)** Quantitative proteomic data retrieved from Nam et al^51^ show that JARID2, DTNBP1 and ASTN2 protein levels are ∼half in KOLF2.1J compared to the reference H9 line at both pluripotency (C) and neural progenitor stage (D). Graphs show mean +/-SD of n=4 cultures and q-values as calculated in the reference proteomic analysis with Welsch’s t-test and FDR correction for multiple comparisons^51^.

**Table 3:** CNVs detected in KOLF2 iPSCs by the analysis of publicly available HumanCoreExome-12_v1 SNP array data.

| Sample | CNVs called in KOLF2.1J | CNV called by PennCNV | CNV called by visual review |
| --- | --- | --- | --- |
| KOLF2 iPSCs | 3p13del | No | Possible mosaic deletion |
|  | 3q14dup | Yes | Yes |
|  | 6p22del | No | No |
|  | 9p22del | No | No |
|  | 18q22dup | Yes | Yes |

Given that the healthy donor does not carry Chr6 and Chr9 CNVs, we could not exclude a deletereus effect. We then reasoned that for a CNVs to cause deleterious effects, it should first alter the expression of the affected genes. Chr6 and Chr9 CNVs cause a complete deletion of *DTNBP1* and partial deletions predicting loss of expression of *JARID2* and *ASTN2.* Indeed, a recent proteomic study shows that compared to H9 hESCs, KOLF2.1J iPSCs express about half of ASTN2, JARID2 and DTNBP1 both at pluripotent and neural progenitor stages (Fig 4C,D)^51^.

Altogether, our work reveals that KOLF2.1J iPSCs carry five CNVs, two of which arose *in vitro* and affect the expression of genes associated with neurological disorders. These findings are essential for the interpretation of neural phenotypes derived from studies using KOLF2.1J iPSCs and to guide the selection of iPSC lines for future studies.

## Discussion

The mutational burden of an hPSC line constitutes a collection of inherited variants and variants arising during the reprograming, passaging, and genome engineering when relevant. Several studies have shown that SNVs and small insertion and deletions (indels) are largely derived from parental cells, whereas large structural alterations (chromosomal rearrangements and CNVs >1Mb) arise during the reprograming or passaging^33–38^. However, less is known about the occurrence of other structural variants (SV), such as smaller CNVs (<1Mb). A typical human genome is estimated to carry 25,000 SVs and a large fraction of these are small structural events which are rarely reported in hPSC genome characterization studies due to the limited resolution of approaches chosen for the analysis^32, 49, 52^. For instance, in the genomic characterization of KOLF2.1J iPSCs only large events (>5Mb) and small indels (<50bp) were confidently assessed^13^. Although the genome sequencing data performed at 30x mean coverage should, in principle, provide the resolution to detect most classes of SVs, there is no single algorithm that can comprehensively call all types of SVs. The algorithm originally chosen for KOLF2.1J genome sequencing analysis depends on the detection of split reads and discordant read-pairs, which are often filtered out due to poor quality indicators or low read coverage. Therefore, a large fraction of KOLF2.1J iPSC genome is likely still uncharacterized.

Nonetheless, even with a complete catalogue of genetic variation, their effects on hPSC performance, differentiation and interaction with other genetic variants for disease modeling would still be difficult to ascertain. Yet, evidence derived from population and disease genomic studies can provide some functional insights. Indeed, mutational constraint metrics indicate that disruptive variants in *JARID2* and *ASTN2* are rare in human genomes thus predicting that KOLF2.1J CNVs that delete *JARID2* and *ASTN2* coding exons are likely deleterious for humans. Accordingly, mounting evidence show that *ASTN2* haploinsufficiency provides risk to neurodevelopmental disorders, and heterozygous loss of *JARID2* is the cause of an autosomal dominant neurodevelopmental syndrome. Furthermore, *JARID2* acts as a hematopoietic tumor suppressor through Cyclin D1 repression and deletions drive transformation to secondary acute myeloid leukemias^53–55^. Together, genetic constraint and disease associations of *JARID2* are reminiscent of *ARID2*, in which a heterozygous 19bp deletion was corrected to generate the KOLF2.1J iPSC line from KOLF2-C1s^13^. Of note, both ARID2 and JARID2 belong to the ARID family, comprised of 15 proteins with diverse functions in development, proliferation and tissue specific gene expression^56^.

JARID2, in particular, is a regulatory subunit of the Polycomb Repressive Complex 2 (PRC2)^57–60^, which maintains transcriptional repression of developmental genes by fostering chromatin condensation through catalyzing histone H3 lysine 27 (H3K27) methylation^45^. Genetic disruption of PRC2 is a major cause of cancer and developmental disorders in humans^61–63^ and postnatal depletion in the mouse brain leads to neurodegeneration due to the de-repression of death promoting genes^64^, while full body disruption causes embryonic lethally^65–67^. Likewise, *Jarid2* deletion leads to gastrulation arrest in *Xenopus* and knock out mice embryos die at embryonic day 10.5-15.5 due to neural tube, heart and hematopoietic defects that vary in severity depending on the genetic background of the mouse strain^58, 68, 69^. Data from mouse ESC (mESC) cultures also support that *in vitro* differentiation is *Jarid2* dose sensitive, given that homozygous and heterozygous *Jarid2* knock out mESCs show delayed differentiation to the three germ layer derivatives, including neurons^57, 59, 70, 71^. In contrast, little is known about the consequences of *JARID2* deletion in hPSCs. The functional characterization of KOLF2.1J iPSCs, which carry only one intact copy of *JARID2,* indicates that global pluripotency and neuronal differentiation phenotypes are comparable to other hPSCs, with the exception of few parameters (i.e. reduced amplitude of evoked excitatory postsynaptic currents in derived neurons)^13^. Furthermore, a recent study shows that the proteome remodeling trajectory during neural induction is similar between KOLF2.1J iPSCs and H9 hESCs, although at steady state level the two lines show profound protein expression differences^51^. Whether KOLF2.1J iPSC-derived neurons display specific neurological disease phenotypes or higher susceptibility to genetic variants associated to neurological disorders remain unknown, but our findings urge to consider these possibilities when working with KOLF2.1J iPSCs.

The identification of CNVs affecting *JARID2*, *DTNBP1* and *ASTN2* may limit the potential of KOLF2.1Js as a “global” reference iPSC line. However, it is likely that such a “global” reference iPSC may never exist. While some users may argue that the phenotypic consequences of carrying at least three potentially deleterious variants are overturned by the overall good growth, editing and differentiation performance of the KOLF2.1J line, users studying neural phenotypes may want to search alternative genetic backgrounds “free” of deleterious variants known to be associated with neurological diseases. This search by itself raises several issues including the lack of a comprehensive catalogue of genomically characterized iPSC lines. Furthermore, although current studies suggest that ∼5% of the apparently healthy human genomes carry a rare loss of function variant known to be deleterious for human development^72^, it is likely that this percentage increases over time as we continue surveying “healthy” and disease associated human genomes with continuously developing sequencing and computational technologies, and therefore no perfect iPSC line exists. While we fill these gaps in knowledge, we propose that the field should work together to generate a catalogue of iPSC lines which includes a comprehensive genome sequencing analysis that enable users with limited resources and computational expertise to interrogate their genes of interest and retrieve unambiguous information about coverage and the type of genetic variants assessed and reported. A similar resource has already been launched for hESCs (https://hscgp.broadinstitute.org/hscgp) and serves as a great example. Ideally, this resource should enable the selection of iPSC lines that do not carry deleterious mutations clearly associated with neurological disorders (or disorders of interest to other studies).

## Limitations of the study

Our study is focused on the five new CNVs incidentally identified by an SNP array in KOLF2.1J iPSCs but our analysis is not comprehensive due to limits in the resolution of the SNP array and targeted analysis applied on the genome sequencing data. Based on estimates for variation in human genomes we expect that KOLF2.1J iPSCs will carry many more SVs that will be revealed with future additional sequencing methods and computational algorithms. Furthermore, while our data show CNVs associated with rare neurodevelopmental disorders in KOLF2.1J iPSCs, the potential of KOLFs to acquire overall characteristic features of neural cell lines upon diverse differentiation protocols has been recapitulated by multiple independent laboratories. Thus, the impact of these CNVs for pluripotency and neural cell phenotypes, as well as their interaction with other variants that researchers may want to introduce to generate experimental models of neurological disorders, remain unknown.

## Acknowledgments

We thank members of the iNDI initiative for their open discussion and encouragement to publish our findings as aligned with their transparency policy. We also thank Dr. Skarnes for kindly sharing with us KOLF2.1J and NGN2-KOLF2.1J iPSCs early on. This work was supported by NIH/NINDS R01NS119699 and NIH/NICHD R21HD107592 (to N.A.), by the CZI Collaborative Pairs Award (to R.A.N. and E.J.B.) and by technical consultation from the IDDRC Biostatistics and Data Science core (HD105354).

## Author contribution

C.G-D., J.E.P., M.E.K., B.D., G.G.C., S.L., A.G., J.A.M., and T.R. conducted experiments. C.G-

D, J.E.P., K.W., J.G. and N.A. designed experiments and wrote the paper. All the authors contributed to data interpretation and edited and reviewed the paper. K.W. supervised the genome sequencing reanalysis. J.G. supervised the SNP array CNV analysis. N.A. supervised and coordinated the project.

## Methods

### hPSC research

iPSCs and ESCs used in this study were previously generated from material obtained under informed consent and appropriate ethical approvals. All the work was performed under approval from the Children’s Hospital of Philadelphia Institutional Review Board (IRB) and Institutional Biosafety Committee (IBC) and following the ISSCR standards for human stem cell use in research.

### hPSC culturing

KOLF2.1J human pluripotent stem cells (JAX, JIPSC1000) were cultured on Matrigel (Corning, #354277) coated plates in mTeSR1 media (STEMCELL Technologies, #85850). The medium was changed every day and cells were passaged when 70% confluent using ReleSR (STEMCELL Technologies, #100-0484) at a 1:20 ratio. When thawing KOLF2.1Js 10 µM Rho-associated protein kinase (ROCK) inhibitor (Y-27632) (Tocris, #1254) was added into the culture media overnight.

### Genomic DNA extraction and quantitative PCR

hPSC pellets were processed for gDNA extraction using the GeneJET Genomic DNA Purification Kit (Thermo Scientific, #K0721). Eluted gDNA was quantified using NanoDrop. gDNA qPCR was performed in a 10 µL reaction volume containing 12ng of gDNA, 0.5 µM of forward primers, 0.5 µM of reverse primers (Supplementary Table) and 1X Power SYBR Green PCR Master Mix (Thermo Fisher, #4367659) in a 384-well plate (Thermo Fisher, #AB1384W) and ran using the CFX384 Touch Real-Time PCR Detection System (Bio-Rad, #1855484). Primers were designed with Primer3^73^.

### Genomic DNA PCR and Sanger Sequencing

Primers targeting sequences upstream and downstream the breakpoints of Chr6 and Chr9 deletions CNVs were designed for PCR and Sanger Sequencing (Supplementary Table). PCR was performed in a 30 µL reaction volume containing 1X Colorless GoTaq Reaction Buffer (Promega, #M792B), 200 µM dNTPs (Invitrogen, #18427013), 0.5 µM forward and 0.5 µM reverse primers, GoTaq DNA Polymerase (Promega, #M3008) and 1µL of template gDNA. To prepare PCR products for Sanger Sequencing we enzymatically purified the products using ExoSAP-IT PCR Product Cleanup Reagent (Applied Biosystems, #78201.1.ML). Purified PCR products were mixed with the forward primer at 1.67 µM and Sanger sequenced in Genewiz.

### SNP genotyping array

Sample gDNA was hybridized to an Illumina Infinium GSA-24v3 BeadChip, consisting of ∼650,000 short invariant 50 mer oligonucleotide probes conjugated to silica beads. After hybridization a single-base, hybridization-dependent extension reaction was performed at the target SNP which label alternate alleles (herein denoted A and B) with different fluorophores. Arrays were loaded onto an iScan System and scanned to extract data. DMAP files enabled the identification of bead locations on the BeadChip and quantification of the signal associated with each bead. Raw fluorescence intensity from the two-color channels was processed into a discrete genotype call (normalized to continuous value 0-1 B-Allele Frequency (BAF)) and the total intensity from both channels (normalized to continuous value with median=0 Log R Ratio (LRR)) at each SNP informed for copy number. The resulting raw Intensity Data files (*.idat) were converted to Genotype Call files (*.gtc) and then into BAF and LRR signal files.

### CNV detection from SNP genotyping array

PennCNV was used as the main CNV detection algorithm. PennCNV calls were filtered to include CNVs with number of SNPs supporting >=20 and length >=100,000 and Segmental Duplication track coverage < 0.5. Related cell line clones CNV calls were compared to ensure consistency in CNV calling. All genomic coordinates are in GRCh37/hg19 human build version. For KOLF2 CNV analysis, genotyping data containing SNP based BAF and LRR values was downloaded from European Nucleotide Archive (HPSI0114i-kolf_2.wec.gtarray.HumanCoreExome-12_v1_0.20141111.genotypes.vcf.gz), which contains data for 511,966.

### Structural variant analysis of genome sequencing

KOLF2.1J genome sequencing data was downloaded from ADWB (https://doi.org/10.34688/KOLF2.1J.2021.12.14). Illumina paired-end whole genome sequencing reads were aligned to the hg19 reference using minimap2^74^ version 2.17-r941 in paired-end short read mode. To validate CNV regions identified from SNP arrays, we calculate coverage log2 ratios and B-allele frequencies. For the log2 ratio, we use an in-house software tool which employs a 10kb non-overlapping sliding window spanning the CNV and calculates the log2 ratio between mean window and mean chromosome coverage. This value represents fold change in coverage, with positive values indicating high coverage with potential duplications, and negative values low coverage with potential deletions. B-allele frequency (BAF) is calculated as the fraction of reads supporting non-reference alleles at SNP locations. In regions with normal diploid copy number, BAF values cluster around 0.5 for heterozygous variants, and cluster around 1 for homozygous variants. An increased dispersion around 0.67 or 0.33 indicates allelic imbalance corresponding to a duplication, and the lack of clustering around 0.5 indicates allele loss corresponding to a deletion. We use DeepVariant^75^ to call SNPs and obtain BAFs. We then use BCFTools view (version 1.9)^76^ to filter SNP locations by variant quality score >30 and read depth >10. Finally, to obtain base-resolution breakpoints for each CNV, we run the DELLY^77^ SV caller on the alignment file and filter results by region, SV type and a minimum length of 100kb to obtain a single SV candidate for each region.

## Supplemental Information

**Supplemental Figure 1:**
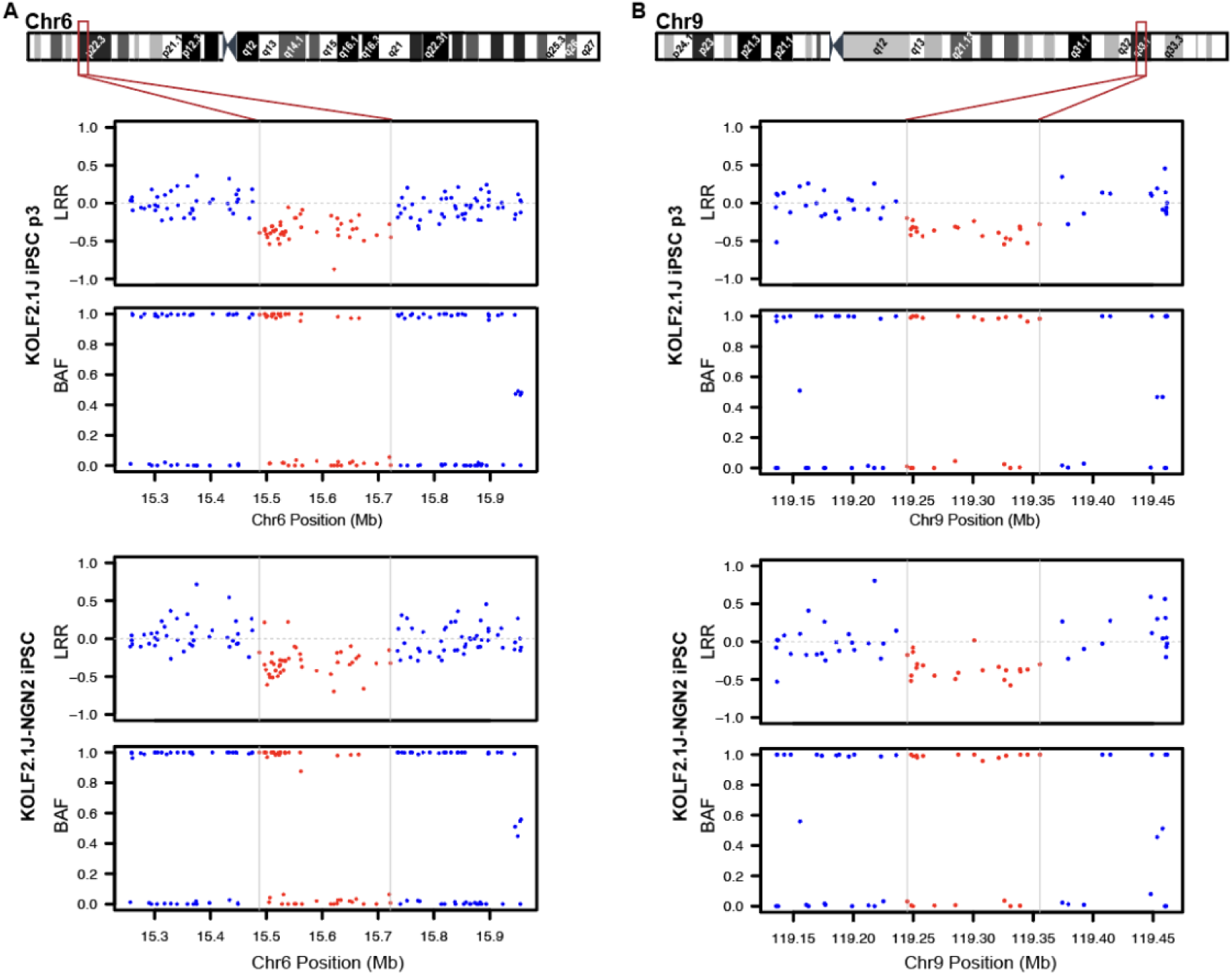
High density SNP array uncovers CNVs affecting coding genes in Chr6p22 and Chr9q33 in a stock of KOLF2.1J iPSCs and KOLF2.1J with doxycycline inducible *NGN2* transgene. **(A)** Chromosome 6 cytoband schematics (top) and Log R Ratio (LRR) and B Allele frequency (BAF) plots (bottom) show reduction of signal intensity and a loss of heterozygosity in 6p22 region of KOLF2.1J iPSCs p3 and KOLF2.1J-NGN2. **(B)** Chromosome 9 cytoband schematics (top) and LRR and BAF plots (bottom) show reduction of signal intensity and a loss of heterozygosity in 9q33 region of KOLF2.1J iPSCs p3 and KOLF2.1J-NGN2.

**Supplemental Figure 2:**
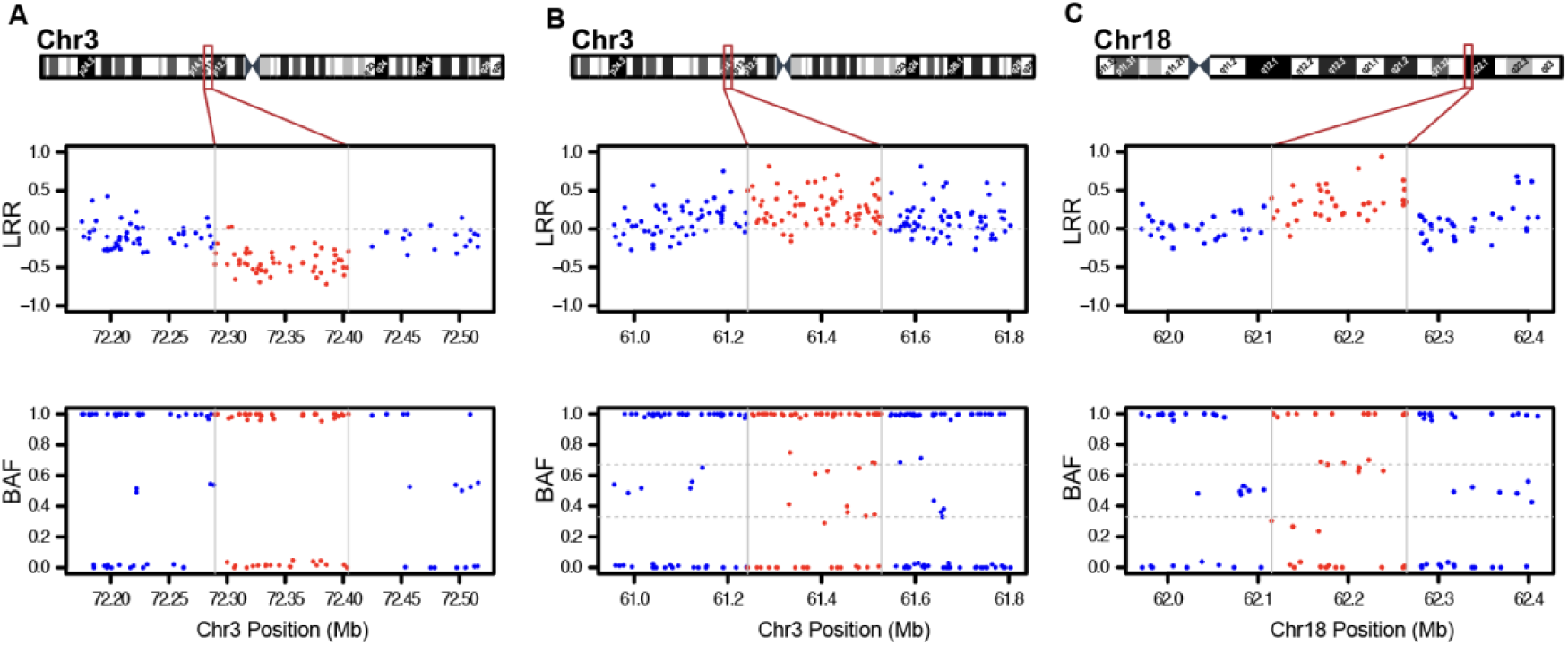
CNVs detected by SNP array in Chr3 and Chr18 do not overlap with coding regions. **(A-B)** Chromosome 3 cytoband schematics (top) and LRR and BAF plots (bottom) showing reduction of signal intensity and a loss of heterozygosity in 3p13 deleted region (A) and increased signal intensity and altered BAF in 3p14 duplicated region (B) of KOLF2.1J iPSCs compared to a control iPSC line. (**C)** Chromosome 18 cytoband schematics (top) and LRR and BAF plots (bottom) showing increased signal intensity and altered BAF in 18q22 duplicated region of KOLF2.1J iPSCs.

**Supplemental Figure 3:**
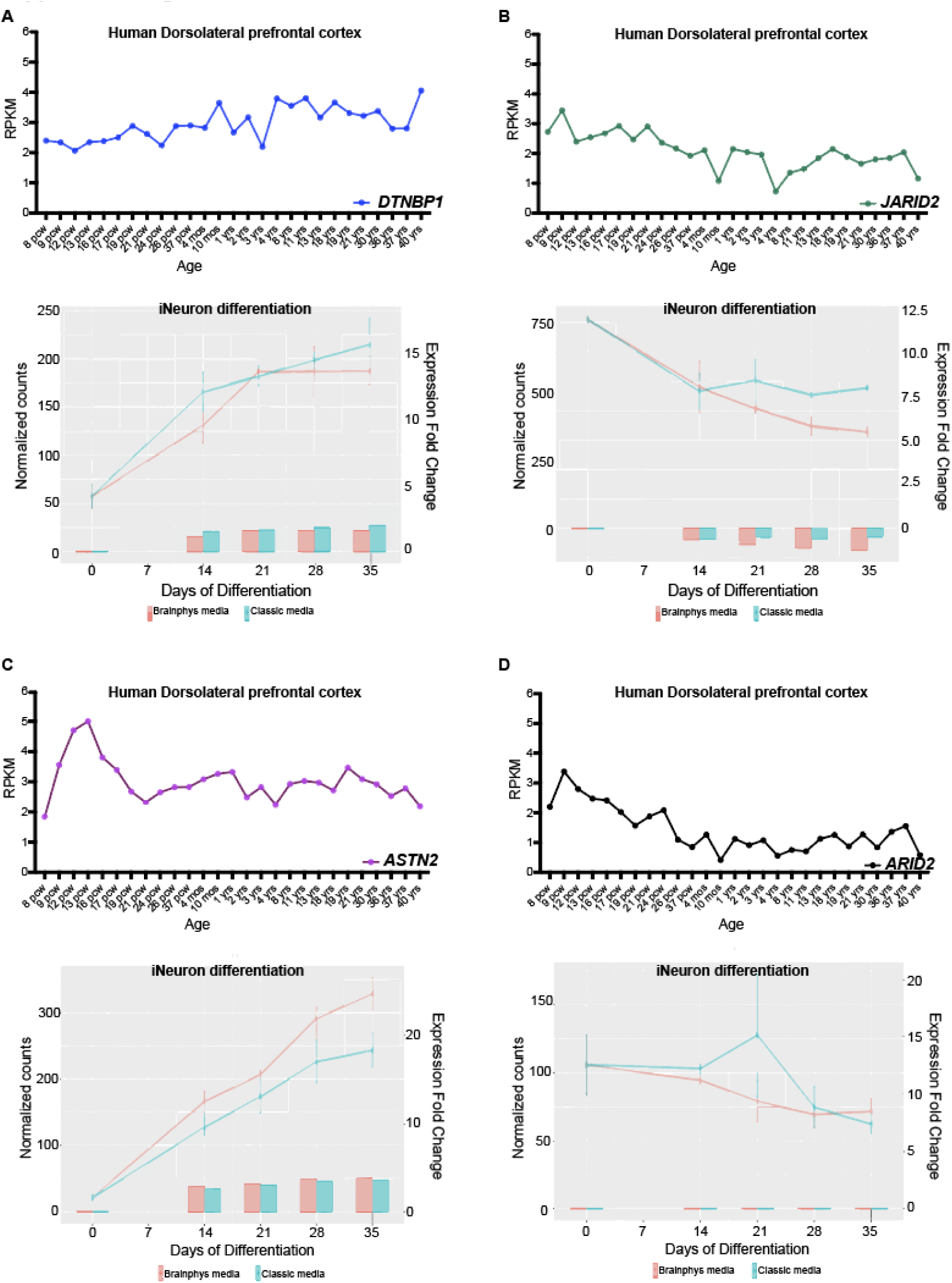
Genes affected by Chr6p22 and Chr9q33 are expressed in neurons in the developing and adult brain. **(A-D)** DTNBP1 (A), JARID2 (B), ASTN2 (C) and ARID2 (D) expression in developing and adult human prefrontal cortex and in hPSC to iNeuron differentiation. Human prefrontal cortex expression data is retrieved from Brainspan (https://www.brainspan.org/rnaseq/search/index.html) and iNeuron differentiation expression plots from the web app created by Connor Ludwig, Kampmann Lab (https://kampmannlab.ucsf.edu/ineuron-rna-seq). Pcw; postconception week. Yrs; years.

**Supplemental Figure 4:**
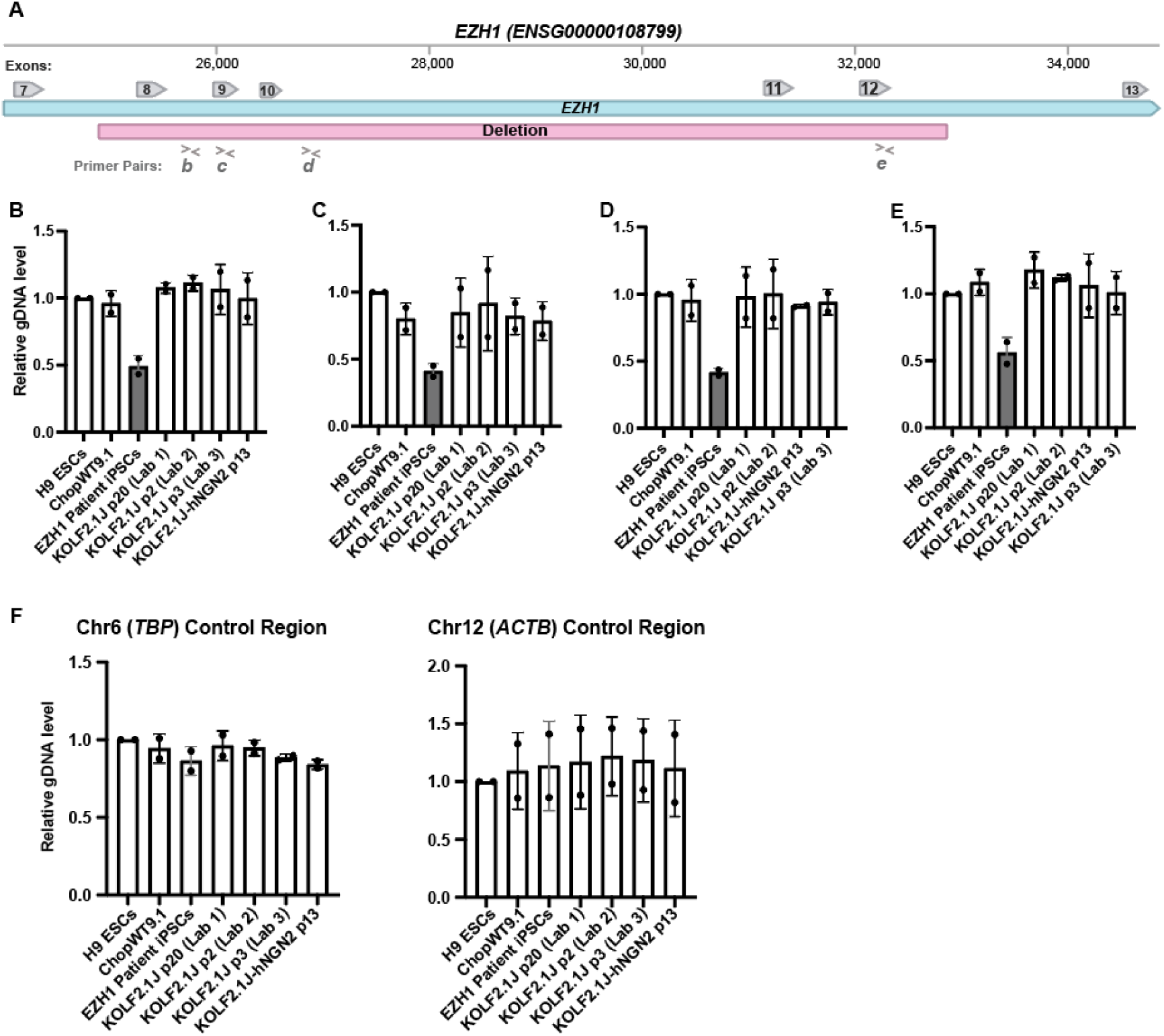
gDNA qPCR on an iPSC line with one copy of EZH1 exon 8-12 deletion demonstrates sensitivity of the technique for detection of hemizygous regions. **(A)** Schematic representation of *EZH1* exon 7-13. Red line illustrates the region deleted in a EZH1 patient iPSC line. Arrowheads (>, <) labeled with b-e indicate position of the primers used for gDNA qPCR. (**B-E)** gDNA qPCR results showing half levels amplification in regions deleted in EZH1 patient iPSCs compared to control and KOLF2.1Js that are expected to be diploid. (**F)** Randomly selected regions of the genome show similar amplification levels across all the hPSC lines.

**Supplemental Figure 5:**
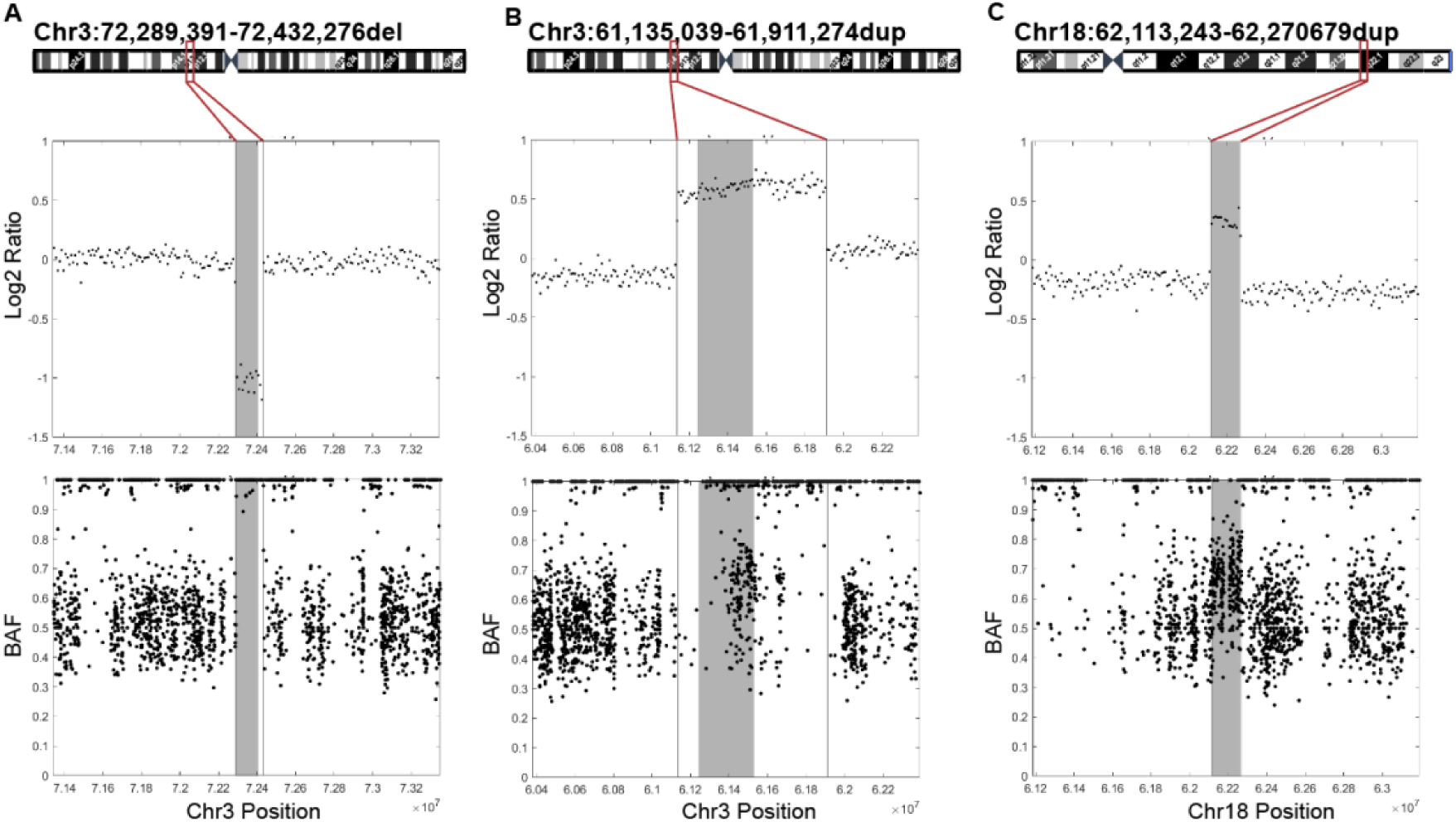
Genome sequencing reanalysis confirms CNVs at Chr3p13, Chr3p14 and Chr18q22. **(A-C).** Chromosome cytoband schematics with base resolution break points of the CNVs (top) and LRR and BAF plots (bottom) obtained from the KOLF2.1J iPSC genome sequencing reanalysis. LRR plots show reduction of signal intensity and BAF plots show loss of heterozygosity in Chr3p13 (A) and gain of signal intensity and altered BAF in Chr3p14 (B) and Chr18q22 (C) CNV regions compared to diploid up and downstream regions. Shadowed area represents the CNV defined by the SNP array and vertical lines the base-resolution breakpoints determined from the genome sequencing data.

**Supplemental Figure 6:**
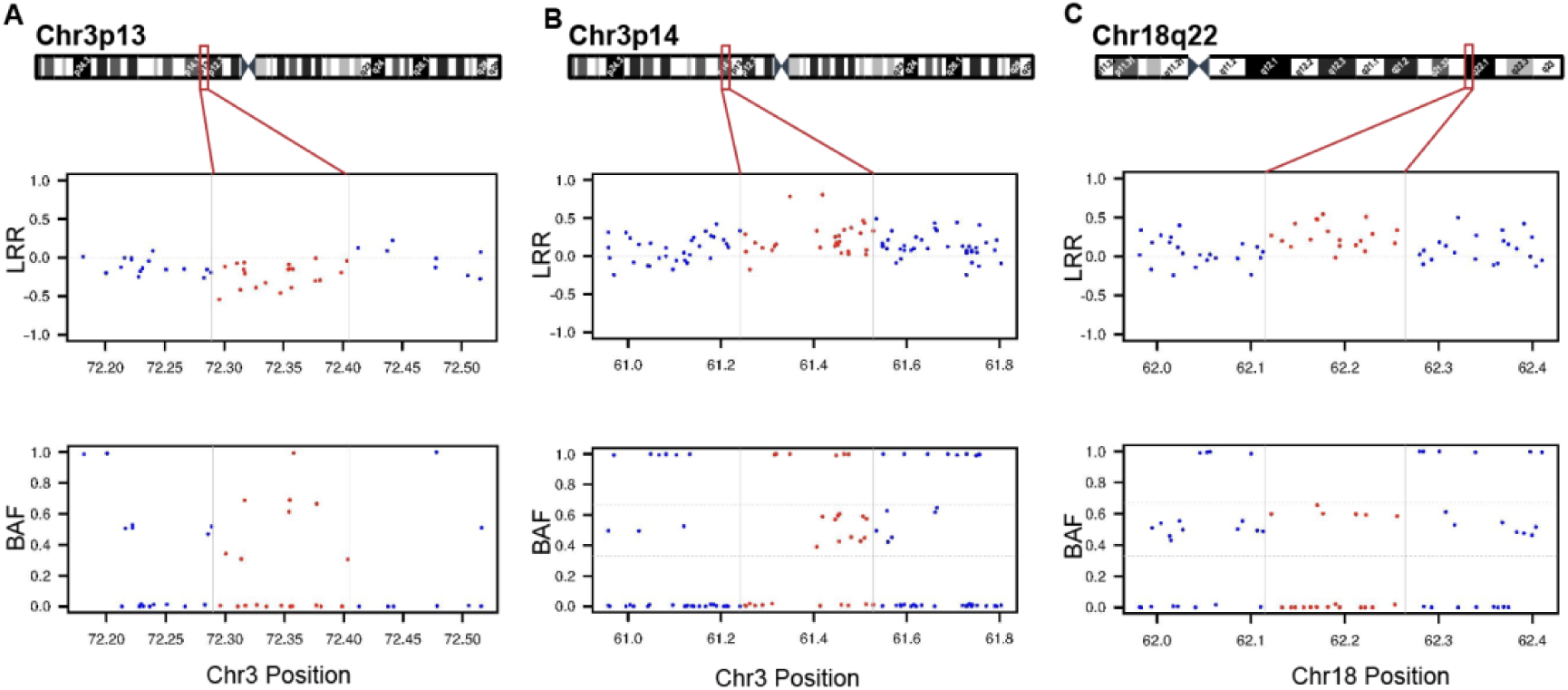
Chr3p13, Chr3p14 and Chr18q22 CNVs in KOLF2.1J were inherited from KOLF2 iPSC line. **(A, B)** LRR and BAF plots of the KOLF2 iPSC line SNP array deposited in HipSci, show that KOLF2 iPSC line carries two of the Chr3p14 (B) and Chr18q22 (C) CNVs in heterozygosity and is likely mosaic for Chr3p13 (A) deletion.

**Supplementary Table 1:**
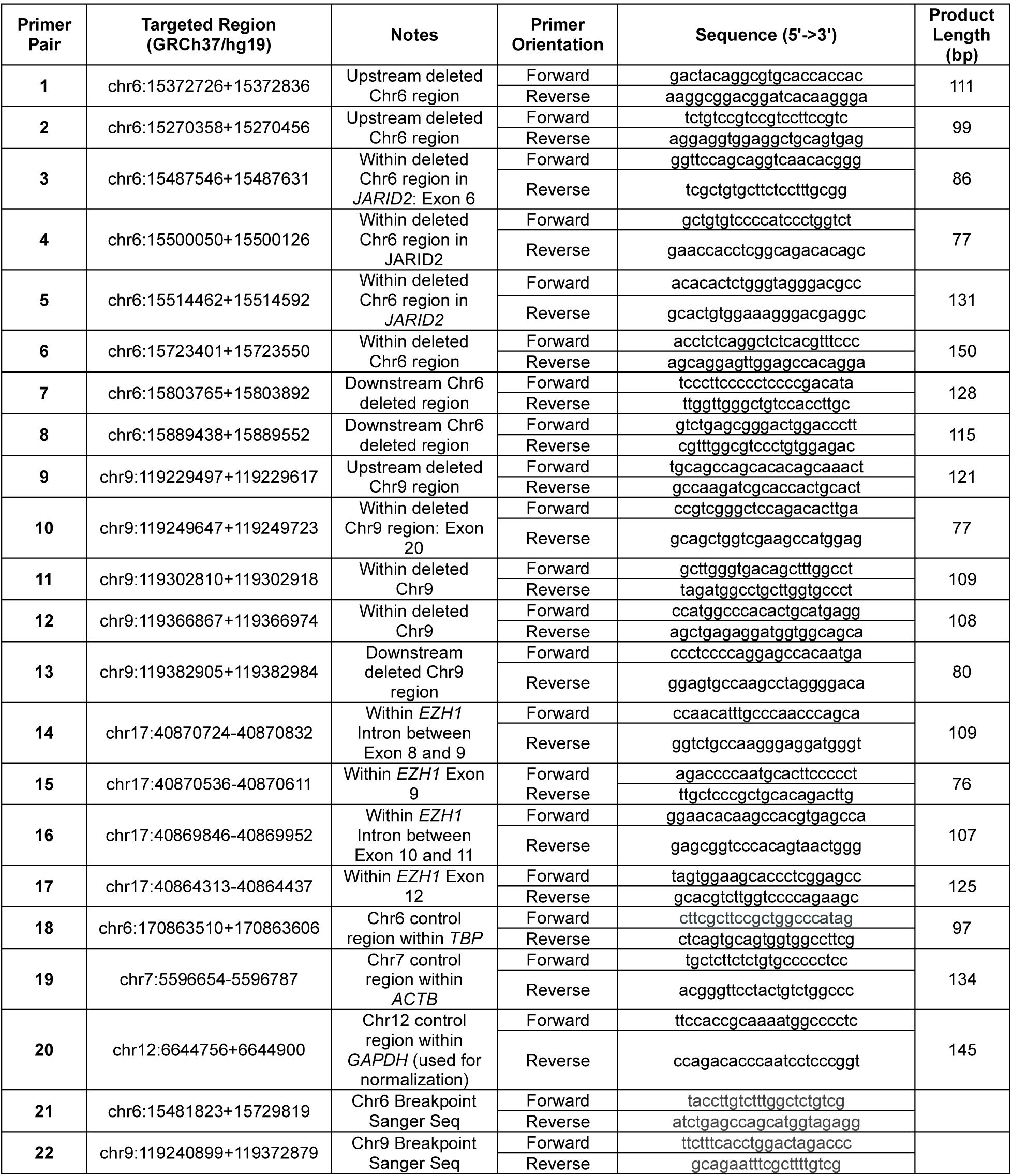
Description and sequence of primers used across the study.

